# Satellite subgenomic particles are key regulators of adeno-associated virus life cycle

**DOI:** 10.1101/2020.10.20.346957

**Authors:** Junping Zhang, Ping Guo, Xiangping Yu, Kiwon Lee, Jenni Firrman, Matthew Chrzanowski, Kuntao Chen, Xiongwen Chen, Derek Pouchnik, Yong Diao, Richard Jude Samulski, Weidong Xiao

**Author notes:** These authors contributed equally to this work (Junping Zhang, Ping Guo, Xiangping Yu).

## Abstract

Historically, AAV defective interfering particles (DI) were known as abnormal virions arising from natural replication and encapsidation errors. Through single virion genome analysis, we revealed that a major category of DI particles contains a double stranded DNA genome in a “snapback” configuration (SBG). The 5’-SBGs include the P5 promoters and partial rep gene sequences. The 3’-SBGs contains the capsid region. The molecular configuration of 5’-SBGs allowed double stranded RNA transcription in their dimer configuration, which in turn regulate AAV rep expression and may improve AAV packaging. In contrast, 3’-SBGs at its dimer configuration increased levels of cap protein. The generation and accumulation of 5’-SBGs and 3’-SBGs appears to be coordinated to balance the viral gene expression level. Therefore, the functions of 5’-SBGs and 3’-SBGs may help maximize the yield of AAV progenies. We postulate that AAV virus population behaved as a colony and utilizes its subgenomic particles to overcome the size limit of viral genome and encodes additional essential functions.

## Introduction

AAV is a replication defective parvovirus which requires a helper virus to complete its life cycle (*1*). The virus is best known for its small genome, which is tightly packaged inside a capsid that is 20-25nm in diameter. Its single stranded DNA genome contains approximately 4700 nucleotides and encodes the rep and cap genes which represent the non-structural and capsid proteins, respectively. AAV replication is mediated by the rep proteins, which are capable of nicking the inverted terminal repeats (ITR) and therefore initiates AAV replication. In the presence of a helper virus, AAV undergoes its lytic infection. Without a helper virus, AAV integrates into the host genome and maintains a relatively stable, latent state.

AAV infection is highly regulated in both latent and lytic infections. In latent infection, AAV genes remain silent. In the lytic life cycle, rep78 and rep68, under control of the P5 promoter, are expressed early, followed by expression of the packaging related genes, such as rep52/rep40 and capsid genes VP1, VP2 and VP3. In the late stage of AAV lytic infection, expression from the P5 promoter is downregulated to facilitate virus encapsidation, which was also demonstrated in recombinant AAV replication and packaging (*2*). A variety of host, viral, and helper virus factors are involved in the temporal and spatial regulation of viral replication and packaging.

Although the canonical AAV particle typically contains both ITRs and the entire coding region, AAV populations are often found to be heterogenous (*3*–*7*). Upon processed through a CsCl gradient, the full AAV particles are present in the fraction with 1.4g/ml density. Lighter density AAV particles (1.32 and 1.35g/ml) are largely deemed as empty particles or defective interfering (DI) particles. DI particles contain aberrant, shorter AAV DNA genomes, and are named for their ability to inhibit AAV replication intracellularly. Template switch was proposed to explain the nonunit-length AAV genome (*7*). However, their composition and molecular conformation are elusive.

In this study, we systemically sequenced the genomes of AAV viruse within a population and revealed the molecular state of AAV genomes, both full length genome and DI particles at the single virus level. Unexpectedly, we discovered that a major class of DI particles are capable of regulating important biological functions for AAV expression and are an integral part of the AAV life cycle. In contrast to the general belief that all subgenomic particles are waste-byproducts and only compete for essential resources required for wild type virus replication and packaging, our findings suggest that the AAV virus has evolved to utilize these subgenomic particles to encode dsRNA that regulate Rep expression and augment Cap expression. This is a clever manipulation evolved by AAV to circumvent its size limitation.

## Results

### Molecular state of AAV subgenome particles at single virus level

The high GC content and palindromic nature of the AAV ITR has been a major obstacle for analyzing AAV genomes. The full DNA genome of the AAV virus has largely been obtained through assembling multiple fragments to obtain a consensus sequence. Even with next generation sequencing, the library construction procedures often require breaking the sample DNA and reassembling the genomes. At the single virus level, important information became lost by this maneuver since the viral genomes in a population are derivatives of the same consensus sequences. Here we utilized the Pacbio Single Molecule, Real-Time (SMRT) sequencing platform and mapped the population genomic configuration distribution of the AAV genome at a single virus level (Fig. 1). There was no enzymatic or mechanic actions that altered the original viral DNA configuration. From analysis of more than 400,000 ccs reads, four major categories of molecules are found in the AAV population. Category 1: Canonical AAV genomes, which contain the full AAV genome flanked by two copies of the AAV ITR; Both positive and negative strands are encapsidated equally into AAV capsids. Category 2: Snapback genomes (SBG), which contains partial duplex AAV genomes with an ITR. Such molecule was shown to be able to snap back and anneal to itself upon denaturing and renaturing cycle (*7*). SBGs are essentially a self-complementary DNA molecule with either the left moiety (5’-SBG) or the right moiety (3’-SBG) genomes. These snapback genomes were either symmetric or asymmetric. The symmetric SBG has near equal length of top strand and bottom strand when in self-complementary conformation. The asymmetric SBG has varying lengths of top strand or bottom, which leads to a single strand region as a loop. Category 3: Incomplete genomes (ICG), which contain an intact 3’ITR and a partial AAV coding sequence, but the sequences do not reach the 5’ ITR. Category 4: Genome deletion mutants (GDM), in which the middle region of the AAV genomes are missing. The molecular distribution of GDM is shown Fig. S1 and the ratio of ICG vs SBG was presented in Fig. S2. Any viral particles that do not have the canonical AAV genomes are referred to as subgenomic particles (SDG), which may include those containing host genomic DNA and helper DNA, which were not presented here in details.

**Fig 1.**
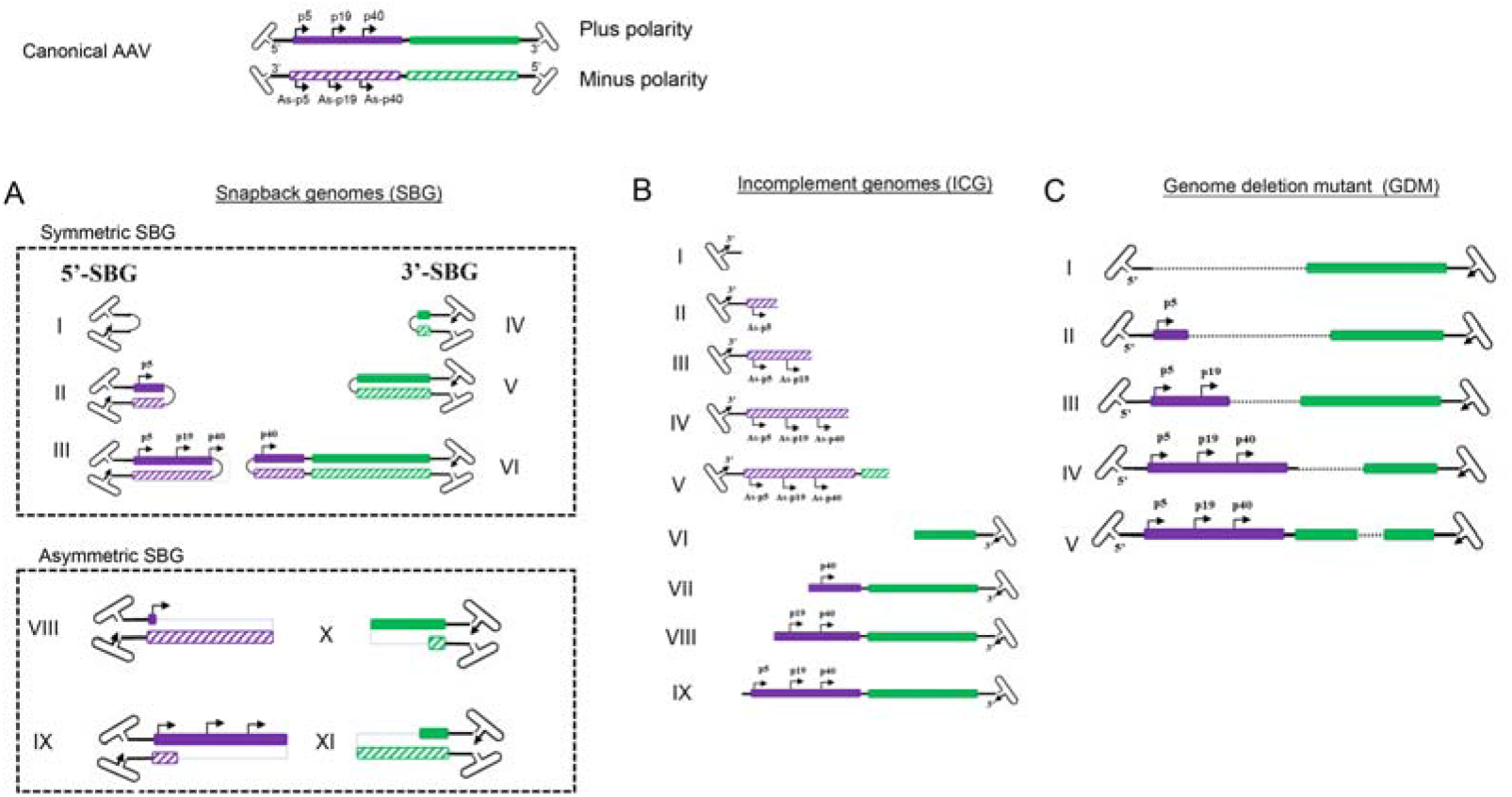
Illustration of molecular configurations for major categories of subgenomic particles in a wild type AAV population summarized from 409919 sequencing reads of two PacBio sequencing runs (220645 and 189274). Both plus and minus strands of the canonical AAV genome are shown at the top. A. Varying lengths of snapback AAV genomes (SBG) are illustrated, including both symmetrical snapback genomes (sSBG) and asymmetrical snapback genomes (aSBG). B. Incomplement genomes (ICG) in the AAV population are missing the 5’ terminal sequences. C. Genome deletion mutants (GDM) have both 5’ and 3’ ITB and miss in the mid-region of AAV. Only plus strand particles are plotted.

### dsRNA molecules are the hallmark of 5’ snapback subgenomes (5’-SBG) which function as a cis-negative regulator of rep expression

5’-SBG assumed a self-complementary configuration and was capable of replicating in the presence of Rep proteins and adenovirus helper functions, which indicated that the dimeric 5’-SBG molecule consisted of the P5 promoter followed by an inverted, head to head coding region. Therefore, logically dsRNA could be expressed from the dimer 5’-SBG when 5’-SBG undergoes DNA replication and becomes a dimer. Such dsDNA overlaps with p5 transcripts in AAV replication which would be able to downregulate P5 promoter expression, mainly Rep78 or Rep 68. To demonstrate the dsRNA effect on rep expression, we engineered a 5’-SBG dimer with a CMV promoter, partial rep sequence was constructed to assume a tail-to-tail dimeric configuration (Fig. 2A). The RNA transcript would therefore assume dsRNA configuration when folding on itself. As shown in Fig. 2A, the expression of Rep78 was indeed, significantly reduced in the presence of dimer 5’-SBG, which confirmed the function of dsRNA against Rep gene as the trans-active regulation of rep expression.

**Fig. 2.**
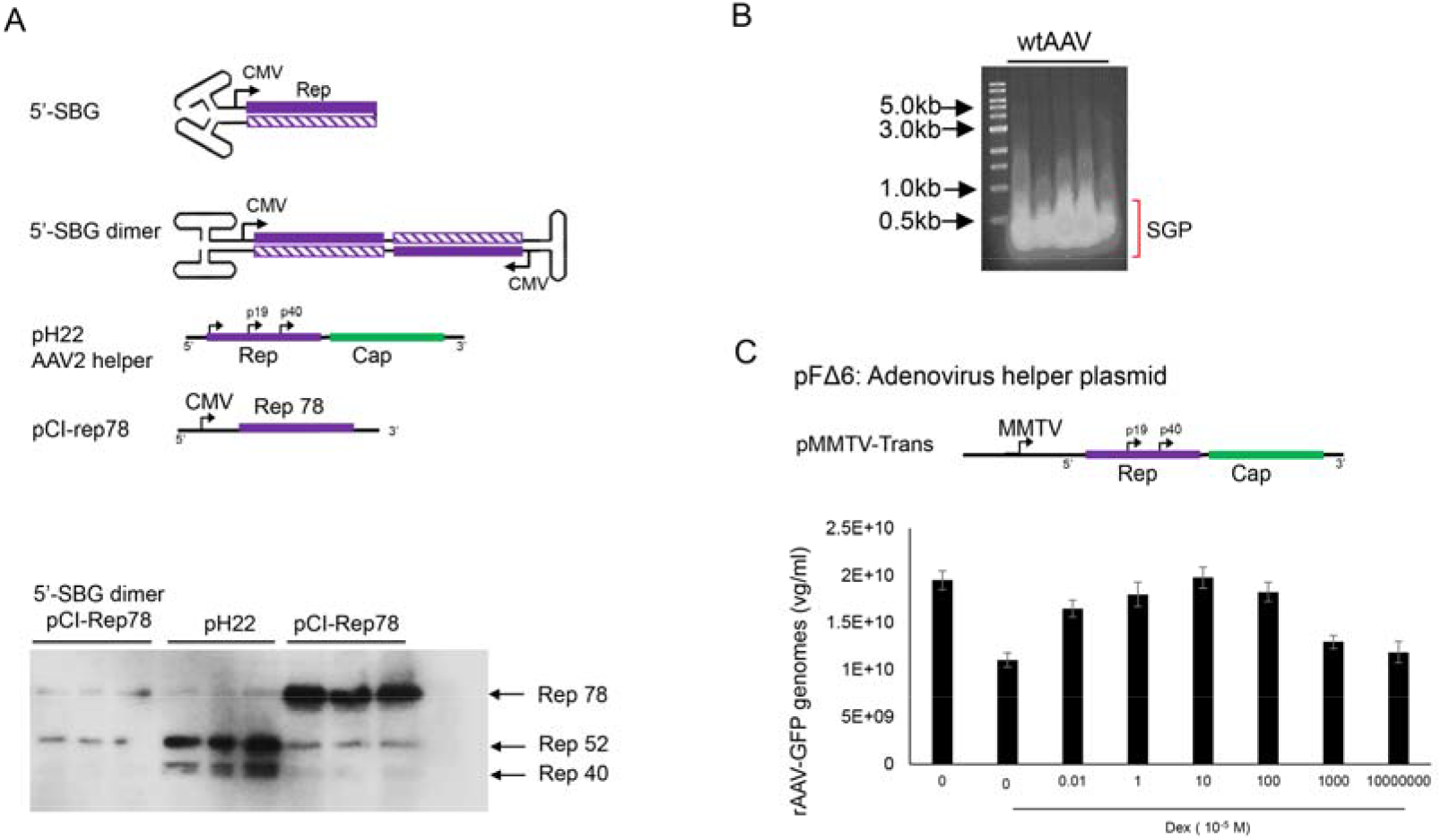
A. Effects of AAV 5’-SBG dimers on rep gene expression. Top panel, sketch of a 5’-SBG in self complementary configuration; 5’-SBG dimer is extended form of 5’-SBG after 2nd stranded DNA synthesis; pH22 is the AAV2 helper plasmid expressing low level of Rep78; pCI-rep78 is the high Rep78 expression plasmid. Bottom panel, Hek 293 cells were transfected with pCI-Rep78 with or without the presence of 5’-SBG dimers. 5’-SBG dimers were made as detailed in the experiment methods. The transfected cells were harvested at 24 hours posttransfection and the expression of proteins were electrophored in 10% PAGE gel. The rep proteins were detected by western blot using mouse anti-AAV-Rep 76.3 antibody. Shown in the figure are triplicate of each testing condition. Effects of AAV subgenomic particles (SGP) on rAAV replication and packaging. B. AAV SGP were separated in an agarose gel and less than 1kb SGP DNA identified in an agarose gel was isolated for the transfection. C. Adenovirus helper plasmid pFΔ6, pMMTV-trans and pssAAV-CB-GFP were co-transfected into 293 cells with or without SGP DNA. Rep expression was under the control of a MMTV promoter in pMMTV-trans, which is inducible by Dexamethasone. rAAV-GFP was collected at 72 hours post-transfection and measured by qPCR. The X axis showed the copy number of SGP molecules used for transfection. “0” means no SGP DNA was added. The cells treated with DEX (10-5M) are indicated in the figure. Y axis: rAAV-GFP yield expressed by vector genomes per ml.

It has been previously shown that down-regulation of Rep 78 gene expression improves rAAV packaging (*2*). To test if 5’-SBG may exhibit such regulatory role, we applied extracted 5’-SBG DNA to a rAAV production system containing pMMTV-trans as the AAV rep and cap expression plasmids (Fig. 2B). MMTV is an inducible promoter which can be activated by dexamethasone. In the absence of dexamethasone, low expression of rep expression gave rise to a higher rAAV yield. Conversely, when dexamethasone was added, the vector yield was reduced. When the 5’-SBG molecules were added to the production system in the presence of dexamethasone, rAAV vector yield was increased at low concentration of 5’-SBG genomes. However, when the amount of 5-SBG was increased to more than 10 copies per cell, it started to exhibit an inhibitory effect, and the vector yield was reduced from the peak. Therefore, it was concluded that 5’-SBG had a regulatory function which senses the AAV genome pool. When SBG/AAV ratio is at a low level, it increases AAV packaging efficacy. However, when SBG is present in excess, it had an inhibitory effect and became a true “defective interfering particles”.

### Bidirectional transcription from 3’ snapback subgenomes (3’-SBG) boosts capsid protein level to maximize vector yields

Similar to 5’-SBG, 3’-SBG replicates in the presence of rep proteins and external AAV helper function. However, the dimeric configuration of 3’-SBG was elucidated as a head to head molecule with the P40 promoter in the center of the dimer. This means that any enhancer in the proximity of the P40 promoter would also affect the neighbor P40 promoter on the opposite strand, which would increase the strength of P40 promoter and expression of the cap protein. To demonstrate the double enhancer effects of 3’-SBG, 3’-SBG monomeric and dimeric constructs were engineered and the expression of the capsid genes analyzed. As shown in Fig. 3, capsid expression from 3’-SBG monomer containing a single P40 promoter was low, whereas expression from 3’-SBG dimer was significantly increased.

**Fig. 3.**
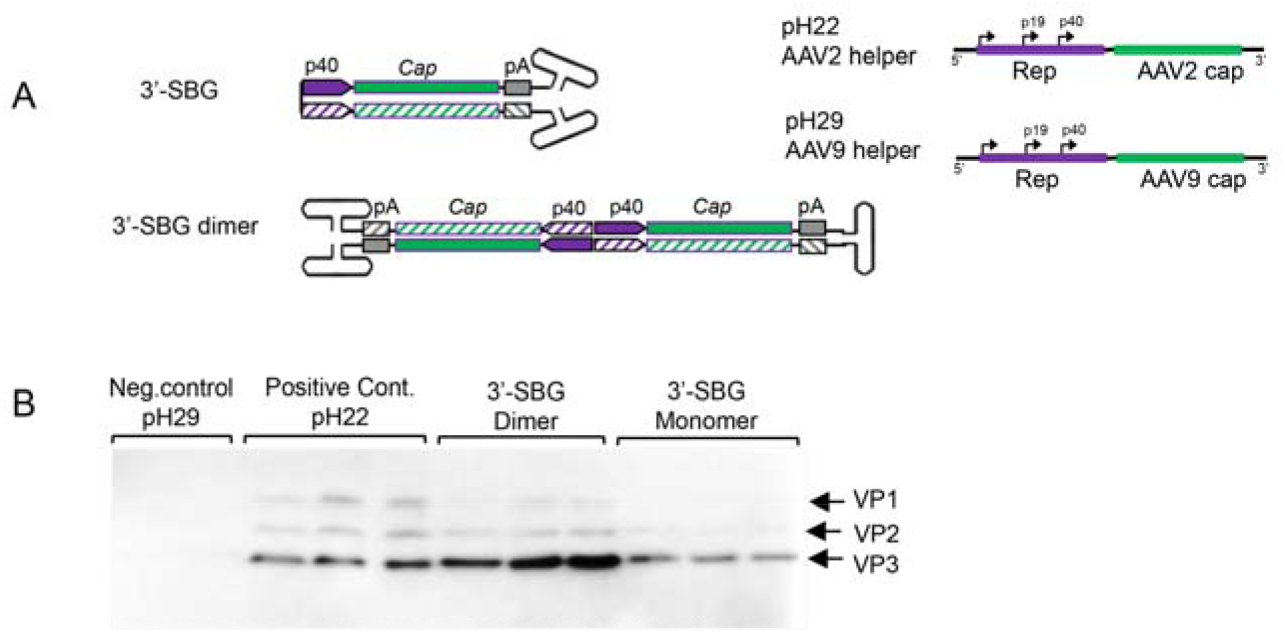
Enhancement of Cap protein expression by 3’-SBG. A. 3’-SBG is shown in monomer configuration; 3’-SBG dimer is the extended form of 3’-SBG after 2nd stranded DNA synthesis or replication. B. Effects of 3’-SBG on capsid protein expression. AAV 3’-SBG molecule dimer was modeled by intermolecular ligation of a SamI and BamHI digestion fragment from the AAV infectious clone psub201. AAV 3’-SBG monomer was modeled by an intramolecular ligation of the same SamI and BamHI digestion fragment. Cap protein expression was detected with mouse anti-AAV Capsid antibody by western blotting at 72 hours post transfection. pH29 negative: HEK 293 cells were transfected with pFΔ6 and pH29 mutation plasmid without AAV2 capsid expression; pH22: HEK 293 cells were transfected with pFΔ6 and pH22 plasmid which express AAV capsid; AAV 3-SBG dimer: HEK 293 cells were transfected with pFΔ6 and 3-SBG dimer molecules; AAV 3-SBG dimer: HEK 293 cells were transfected with pFΔ6 and 3-SBG monomer molecules. Each condition is shown as triplicate.

## Discussion

Previously, the heterogeneity of AAV genomes within a population was not fully characterized because the palindromic structure of the AAV ITR inhibited progression of the typical sequencing reaction, thus requiring AAV genome to be broken/interrupted in order to make compatible libraries that were compatible for the sequencing instrument. However, this maneuver causes the loss of the detailed information of individual molecules, since DNA in AAV subgenomic particles are inherently diverse and heavily rearranged from the standard genomes (Fig. 1). Reported studies using PacBio to sequence scAAV only recovered special categories of AAV molecules and did not provide details on the individual genomes within the entire AAV population (*8*), In studies with single stranded DNA genomes (*9*), only annealed AAV genomes were captured in the library. Here we have obtained the full profile of AAV genomes within a population at the single virus level (Fig. 1). Those unannealed molecules are also fully sequenced. This high-resolution analysis allowed us to identify and characterize an array of genome configurations that are present. Besides canonical AAV full length genomes, the presence of particles with incomplete AAV genomes, starting from the 3’ITR that failed to reach the 5’-ITR, which indicated an aborted packaging process (Fig. 1). We did not observe obvious hot spots in ICG population, which suggested that incomplete packaging of the AAV genome was most likely a random event.

In addition, we revealed the divergence of genome deletion mutants (GDM) in the AAV population supplementary data Fig. S1. The existence of GDMs as well as the discovery of asymmetric SBG configuration are the primary reason that we proposed that NHEJ is the mechanism of subgenomic particles formation in the AAV population (citation of Preprint). NHEJ is the mechanism that can simultaneously explain the formation sSBG, aSBG, GDM and various forms of subgenomic particles identified in the rAAV population (including those containing foreign genetic element such as host genome DNA and helper genes). Although we did not study GDMs, some GDMs will bring AAP reading frame closer to either P5, 19 or P40 promoter, which would increase AAP expression and enhance AAV package. This possibility is outlined in Fig. 4.

**Fig. 4.**
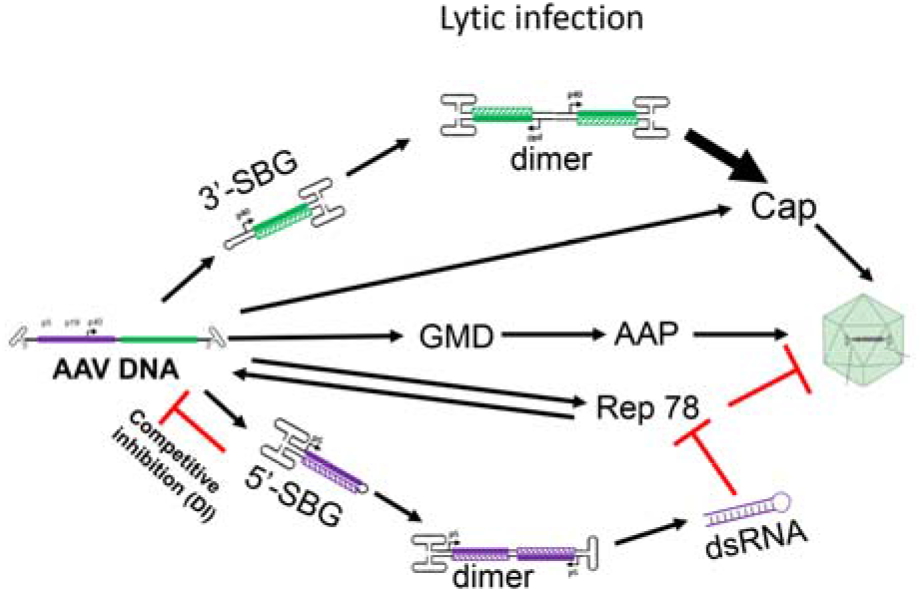
A model summarizing the role of 5’-SBG and 3’-SBG in the AAV life cycle. 3’-SBG improves AAV cap expression when its copy number is increased along with its double enhancer effects in dimer conformation. 5’-SBG expresses dsRNA against Rep78. Rep78 in excess is detrimental to AAV DNA replication and package. Therefore, the relative abundance of AAV genomes and 3’-SBG and 5’-SBG form a dedicated positive and negative loop that ensure an optimum level of production of the next generation progeny. This model is essential since there is dramatic amplification of AAV genomes during replication. The excessive DNA template therefore requires another layer of gene regulation.

AAV SBG molecule is naturally occurring self-complementary DNA genome which is similar to the well-known recombinant scAAV vector which used successfully in clinical trial but does not need a special mutated AAV ITR. In this study, our findings revealed a critical role for SBGs in the AAV life cycle. The 5’-SBG containing the p5 promoter was capable of expressing dsRNA overlapping the rep gene transcript. dsRNA is the precursor for RNAi Interference, which would function as an inhibitor for the p5 promoter transcripts. This is a very clever mechanism that can balance P5 expression levels when the viral template increases excessively at the end of viral replication and packaging and there is no additional need to recruit host factors downregulate the promoter. Since the expression of Rep78 and Rep68 and its inhibitor are under the same transcription mechanism, the inhibition rate is determined by relative copy numbers of full length genome and 5’-SBG, 5’-SBG therefore becomes a de facto AAV genome population sensor. In a hypothetical replication model starting with one full length particle, the p5 promoter initiates expression of rep 78 and rep 68 which replicate the AAV genome. The SBG molecules are generated when AAV genomes have accumulated to a certain level. Since SBG particle replicate faster than the full AAV genome, at a certain point, SBG replication will overtake the full AAV genome and rep78 expression levels will be significantly reduced. Thus, 5-SBG is a rather effective inhibitor and can regulate the AAV genome population. The dsRNA expression was also supported by a previous study which demonstrated the presence of negative stranded DNA extended to the ITR region. 5’-SBG model theory and offer seamless interpretation of those observations (*10*).

One the other hand, the snapback molecules were also formed at 3’-end. 3’-SBG cannot produce dsRNA because the P40 promoters are sitting in a head to head configuration. This leads to the enhancement of P40 gene expression, since the expression can be increased greatly as the copy number of 3’-SBG increases (Fig. 3).

### A model for the essential role of snapback subgenomes in the AAV life cycle

Based on the detailed genomic state and molecular function of the AAV subgenomes, we propose that such molecules are not waste byproduct but play an active and critical role in the AAV life cycle. During replication and packaging, these molecules would function as “cis” and “trans” regulators of rep and cap gene expression, balancing viral gene copy number and expression levels. Without SBG molecules, AAV is primarily replicating itself with reduced packaging. However, the explosive replication of AAV genomes also lead to the production of SBG molecules, which have a growth advantage over the wt AAV genome, and leads to a decrease in rep 78 expression and an increase in cap expression. The end result is an increased packaging of the AAV progenies. This model is summarized in Fig. 4.

The excessive amounts of 5-SBG in the wild type AAV population may have another implication. Based on the results of various genomic studies, we know that during a latent infection of host cells only AAV fragments are found. It is likely that 5’-SBG also functions in the host cells as a suppressor of p5 expression, which may stabilize its latent infection (Fig S3). That may be a reason that it is beneficial to have subgenomic particles present in large quantities when the AAV virus is preparing its latent infection. Protein kinase R (PKR) is activated by dsRNA produced during virus replication. Adenovirus virus-associated RNA-I (VAI) is a short, noncoding transcript that functions as an RNA decoy to sequester PKR in an inactive state. VAI is an essential gene for rescuing AAV from latent infection. It further implies the complicated relationship between AAV helper virus and AAV replication.

AAV virus has a small genome of 4.7k nucleotide in size. Yet its genome design is a rather efficient and space conscience, with all genes overlapping each other in the same reading frame. Nevertheless, the virus developed a mechanism to use subgenomic particles, i.e. defective interfering particles, to express the negative regulator dsRNA, and to enhance expression of capsid proteins to facilitate packaging during the late stage of infection. The presence of such 5’-SBG and 3’-SBG is not co-incidence but evolved over virus spread in human population. The utilization of snap-back molecules is a perfect example of how extra-genomic molecules can be an integral part of AAV regulation. This is the first report describing a virus utilizing these mechanisms, which seems to be shared by all members of the parvovirus family (data not shown). Although these molecules are necessary for the wild AAV life cycle, their presence in the context of rAAV gene therapy is a major concern for long term and stable transgene expression, as well as vector safety.

## Supplementary Materials

Materials and Methods

Fig S1 – S3

Figure legend

